# IgE-mediated regulation of IL-10 and type I interferon enhances rhinovirus-induced Th2 priming by primary human monocytes

**DOI:** 10.1101/432815

**Authors:** Regina K. Rowe, David M. Pyle, J. David Farrar, Michelle A. Gill

## Abstract

Rhinovirus infections are linked to the development and exacerbation of allergic diseases including allergic asthma. IgE, another contributor to atopic disease pathogenesis, has been shown to regulate dendritic cell antiviral functions and influence T cell priming by monocytes. We previously demonstrated that IgE-mediated stimulation of monocytes alters multiple cellular functions including cytokine secretion, phagocytosis, and influenza-induced Th1 priming. In this study, we investigate the effects of IgE-mediated allergic stimulation on monocyte-driven, RV-induced T cell priming utilizing primary human monocyte-T cell co-cultures. We demonstrate that IgE crosslinking of RV-exposed monocytes enhances monocyte-driven Th2 priming. This increase in RV-induced Th2 differentiation was regulated by IgE-mediated inhibition of type I interferon and induction of IL-10. These findings suggest an additional mechanism by which two clinically significant risk factors for allergic disease exacerbations – IgE-mediated stimulation and rhinovirus infection, may synergistically promote Th2 differentiation and allergic inflammation.

## Introduction

The link between viruses and allergic diseases has long been appreciated, and is best represented by the coinciding increase in allergic asthma exacerbations with peaks in fall respiratory virus epidemics (1). Rhinoviruses (RV) are closely associated with allergic disease, representing the most frequently detected viruses in children with asthma exacerbations (1, 2). Furthermore, early RV infection in allergic children also increases the risk of asthma development by up to 8-fold (3). The degree of atopy is similarly linked to disease severity; for example, serum allergen-specific IgE levels in allergic asthmatics directly correlate with virus-induced exacerbation severity (4). Clinically, decreasing serum IgE with omalizumab therapy reduces atopic disease exacerbations (5–7). In a recent study, children with allergic asthma treated with omalizumab had decreased RV-induced upper respiratory symptoms, shortened duration of virus shedding, and decreased peak virus titers (8), suggesting that IgE-mediated processes modulate RV infection *in vivo*.

Monocytes are antigen presenting cells (APCs) recruited to the airway during respiratory virus infections (9, 10) and following allergen challenge (11), whose functions are modulated by the allergic environment (12–15). Several studies indicate roles for monocytes and monocyte-derived cells in RV-induced allergic disease pathogenesis. In asthmatic subjects, experimental RV-infection resulted in exaggerated recruitment of monocytes/macrophages to the airways compared with control subjects which correlated with virus load (9). Our group has shown that IgE crosslinking induces monocyte secretion of regulatory and pro-inflammatory cytokines, including IL-10, IL-6, and tumor necrosis factor (TNF) α (12, 13). IL-10 has been specifically linked to allergic disease pathogenesis and virus-induced wheezing (16–18), although its role in IgE-mediated monocyte antiviral responses remains unexplored. We recently demonstrated that critical monocyte antiviral functions, including influenza-induced upregulation of antigen-presenting molecules and CD4 Th1 T cell priming are inhibited by IgE crosslinking (12). IgE-mediated regulation of monocyte functions during RV infection may thus play a role in promoting allergic inflammation.

Th2-mediated responses, a hallmark of allergic disease, are implicated in RV-induced inflammation. Enhanced Th2 inflammatory responses correlate with increased disease severity in allergic asthmatics experimentally infected with RV (19). While multiple cell types and mediators are likely involved, many studies support roles for both APCs and IgE in promoting Th2 responses. In cat allergic subjects, omalizumab treatment decreased *ex vivo* allergen-induced pDC-driven Th2 stimulation (20); similarly, monocyte-derived DCs from allergen-exposed atopic subjects induced CD4 T cell Th2 cytokine responses (21). These studies suggest that IgE-mediated APC stimulation may regulate Th2 priming during RV-induced allergic inflammation, although the mechanisms underlying this phenomenon remain undefined.

Type I interferons (IFN) are key mediators of antiviral responses and also negatively regulate human Th2 development and effector functions *in vitro* (22, 23). IgE-mediated inhibition of virus-induced APC IFN secretion is one proposed mechanism involved in RV-associated allergic exacerbations (5, 24, 25). IgE crosslinking inhibits influenza- and RV-induced IFN secretion from plasmacytoid dendritic cells (pDCs) and PBMCs (24–26), and omalizumab treatment in children with allergic asthma has recently been shown to restore *ex vivo* antiviral IFN responses (5, 25). Within this study, the group with the greatest post-omalizumab increases in IFN had fewer asthma exacerbations (5), suggesting that virus-induced type I IFN plays an important role in regulating allergic inflammation *in vivo*.

While IgE-mediated effects likely contribute to virus-induced allergic disease, how IgE regulates antiviral responses to RV and promotes Th2-mediated allergic inflammation is unknown. Given the critical role of monocytes and monocyte-derived cells in antigen presentation and T cell differentiation, we investigated the impact of IgE-mediated stimulation on RV-induced monocyte-driven T cell differentiation. We show that IgE-driven effects on monocytes, including inhibition of virus-induced IFN and induction of IL-10, enhance RV-induced Th2 development. These data provide evidence that IgE-mediated alteration of monocyte antiviral functions during RV infection may promote Th2 priming and enhance allergic inflammation.

## Results

### IgE-mediated allergic stimulation enhances RV-induced monocyte-driven CD4 Th2 cell development

We previously established a model of monocyte-driven T cell development to evaluate the effects of IgE-mediated allergic stimulation on monocyte-driven T cell priming (12). Using this system, we determined how IgE-mediated stimulation impacted RV-induced naïve CD4 T cell priming by primary human monocytes (Figure 1 and Supplemental Figures 1 and 2). RV-exposure alone induced an increase in monocyte-driven Th2 (Fig. 1A) and Th1 (Supp. Fig. 2A) differentiation. More notable was the impact of IgE crosslinking plus RV exposure, which significantly enhanced Th2 priming (Fig. 1A and Supp. Fig 1). The percentage of Th2 priming induced by the combination of IgE crosslinking with RV positively correlated with baseline monocyte surface expression of the high affinity IgE receptor (FcεRIα); no correlation was observed with RV exposure alone (Fig. 1B). Similar to our previous findings with influenza (12), the strong Th1 response induced by RV-exposed monocytes was inhibited by IgE crosslinking (Supp. Fig. 1 and 2A). The IgE-mediated enhancement of Th2 and inhibition in Th1 differentiation significantly reduced the Th1/Th2 ratio (Fig. 1C). RV-driven T cell proliferation was also decreased by IgE crosslinking (Fig. 1D). To investigate whether IgE crosslinking similarly impacts virus-specific Th2 priming we co-cultured RV-exposed monocytes with autologous naïve CD4 T cells, followed by restimulation with irradiated RV-exposed autologous PBMCs. Consistent with our allogeneic co-culture results, the combined exposure of monocytes to IgE crosslinking and RV resulted in enhanced CD4 T cell Th2 development (increased IL-4 expressing T cells; Fig. 1E), providing evidence of IgE-mediated effects on both antigen-specific and non-specific Th2 development.

**Figure 1.**
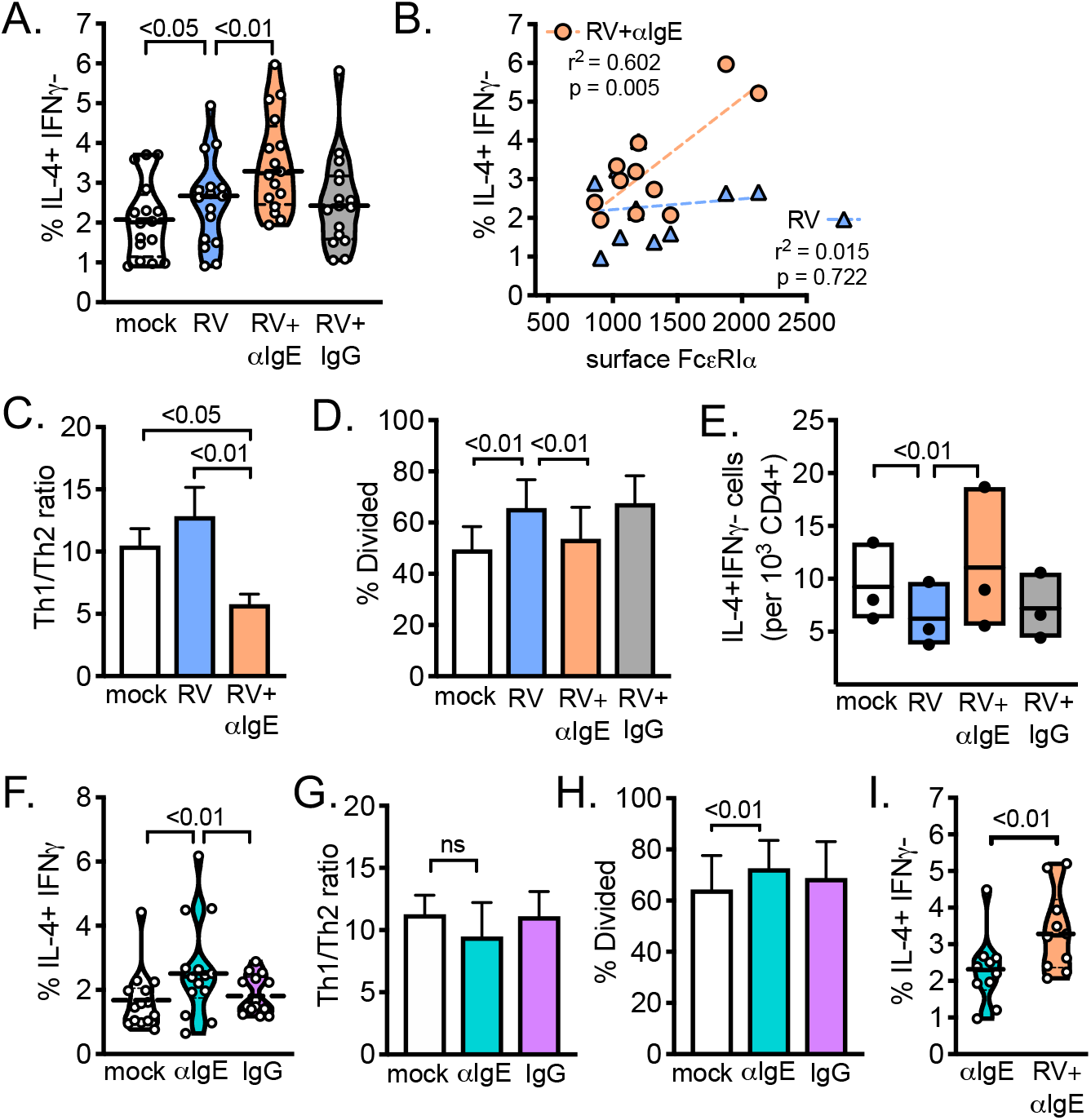
IgE crosslinking enhances RV-induced monocyte driven Th2 differentiation. Primary human monocytes were either mock-(media alone) or RV-exposed in the presence or absence of IgE crosslinking (αIgE) or isotype control (IgG) antibodies, then co-cultured with naïve CD4 T cells to evaluate monocyte-induced Th2 (IL-4+IFN*γ*-) differentiation. (A) RV-induced monocyte-driven Th2 development was measured in an allogeneic co-culture system, n=17 donor pairs. (B) Linear regression analysis between monocyte surface FcεR1α expression and Th2 priming by RV (blue triangles) or RV + αIgE crosslinked (orange circles) monocytes. Pearson correlation coefficient and two-tailed p value were determined, n=11 experiments. (C) Th1/Th2 ratios for donor pairs shown in (A). (D) % T cell division in a subset of experiments, n=10. (E) RV-specific Th2 priming of autologous naïve CD4 T cells, expressed as IL-4+IFN*γ*-cells per 1000 CD4+ T cell events, box plots showing 95% confidence intervals and mean for n=3 experiments, p values by one-way ANOVA with post-hoc comparison. (F-H) Monocyte-driven allogeneic T cell priming in the absence of virus; (F) Th2 priming with the respective (G) Th1/Th2 ratios for n=15 experiments, and (H) % T cell division measured for n=5 experiments. (I) For matched donor pairs in (A) and (F) Th2 priming was compared between IgE crosslinking in the presence or absence of RV, n=10 experiments, student’s 2-tailed paired t-test. All violin plots show the mean depicted as black line with individual experiments as open circles. Bar graphs show mean with standard error. Unless otherwise noted above, p values obtained by two-way ANOVA with post-hoc comparison.

We then investigated the effect of IgE crosslinking on monocyte-driven Th priming in the absence of virus-exposure. Although IgE crosslinking promoted a small increase in monocyte-driven Th2 priming (Fig. 1F), there was no significant change in the final Th1/Th2 ratio (Fig. 1G). In contrast to IgE-mediated effects on virus-driven T cell priming, we observed a small increase in both IgE-mediated allogeneic T cell division (Fig. 1H) and Th1 priming (Supp. Fig. 2B) by monocytes. In the subset of experiments with sufficient cell numbers to directly compare IgE-mediated Th2 priming in the presence or absence of RV exposure, monocytes exposed to IgE crosslinking plus RV promoted greater Th2 priming than monocytes exposed to IgE crosslinking alone (Fig. 1I). These data show that IgE-mediated allergic stimulation combined with RV-exposure significantly enhances monocyte-driven Th2 differentiation, shifting the Th1/Th2 balance towards allergic inflammation. This effect is related to monocyte IgE receptor (FcεRIα) expression, supporting a role for both the virus and IgE in monocyte-driven Th2 differentiation.

### Type I interferon (IFN) negatively regulates RV-induced Th2 development

Our data suggest an interaction between IgE crosslinking and monocyte antiviral responses. To investigate possible mechanisms underlying IgE-induced RV-driven Th2 development, we evaluated the role of type I IFN, a known negative regulator of human Th2 development (23). IgE crosslinking significantly impaired RV-induced monocyte IFNα secretion (Figure 2A). To specifically evaluate the effect of impaired IFN on Th2 priming, we utilized αIFNAR to block type I IFN receptor signaling and found increased RV-induced monocyte-driven Th2 priming in the presence of αIFNAR (Fig. 2B). Consistent with previous findings (23), exogenous recombinant human IFNα reduced monocyte-driven Th2 differentiation in the absence of virus exposure (Supp. Fig. 3A), and conversely enhanced Th1 priming (27, 28) (Supp Fig. 3B) of allogeneic naïve CD4 T cells. Similar to the effects on mock-infected monocytes (Supp Fig. 3A), addition of IFNα also decreased RV-driven Th2 priming (Fig. 2C). Finally, addition of IFNα reversed IgE-mediated enhancement of RV-induced Th2 priming (Fig. 2C), and partially restored IgE-mediated inhibition of Th1 priming (Fig. 2D). These data indicate that IgE-mediated inhibition of RV-induced type I IFN regulates monocyte-driven Th2 and Th1 development.

**Figure 2.**
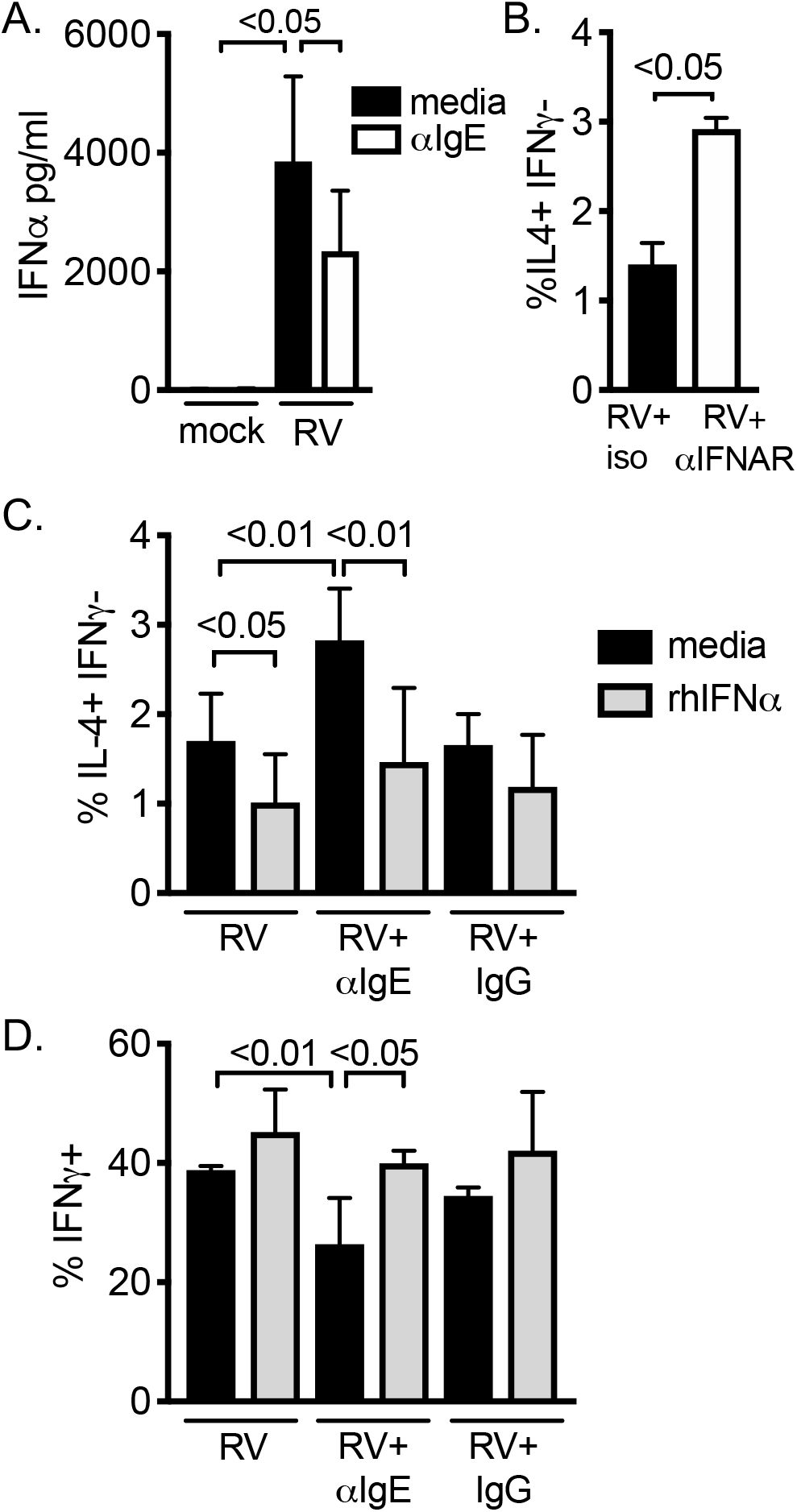
Type I interferon reverses IgE-mediated enhancement of RV-induced Th2 priming by primary monocytes. Monocytes were exposed to either mock or RV conditions in the presence or absence of IgE crosslinking (αIgE) or isotype control (IgG) antibody. (A) Monocyte supernatants were harvested for IFNα analysis at 24 h post infection, n=3 experiments. (B-D) 24h post virus exposure, monocytes were co-cultured with allogeneic naïve CD4 T cells. At the time of co-culture either (B) anti-human IFNAR (αIFNAR) or isotype control antibodies (10μg/mL) or (C-D) rhIFNα (10,000 IU/mL) were added and (B-C) Th2 (%IL-4+IFN*γ*-) or (D) Th1 (%IFN*γ*+) differentiation was measured. Mean with standard error shown, p values by two-way ANOVA with post-hoc comparison, n=3 experiments.

### IgE-induced IL-10 regulates RV-induced Th2 development

Our findings also indicated an IgE-specific factor involved in monocyte-driven Th2 development given the observed increase in the Th2 population by IgE crosslinking both alone and combined with RV (Fig. 1). IgE-mediated stimulation of monocytes induces the production of IL-10 (13, 15), a cytokine known to regulate pro-inflammatory processes including T cell development (29–31). Confirming prior observations, monocytes secreted high concentrations of IL-10 in response to IgE crosslinking, and this was not altered by RV-exposure (Fig. 3A). Recombinant human IL-10 inhibited RV-induced IFNα secretion (Fig. 3B) in monocyte cultures. Addition of exogenous rhIL-10 to monocyte-T cell cocultures also enhanced monocyte-driven Th2 development in the presence or absence of virus (Fig. 3C), and suppressed RV-driven Th1 priming (Fig. 3D). Blocking IL-10 and its receptor (IL-10Rα) reversed IgE-induced enhancement of RV-driven Th2 differentiation (Fig. 3E), without affecting Th1 differentiation (Fig. 3F). Blocking IL-10 in monocyte-T cell co-cultures partially reversed IgE-mediated inhibition of IFNα (Fig. 3G), demonstrating that IL-10 contributes to the IgE-induced negative regulation of RV-induced IFN production in primary monocytes. These results indicate that IgE-mediated IL-10 secretion combined with additional IgE-mediated effects on monocyte functions regulate RV-induced type I IFN to modulate RV-induced Th2 cell development.

**Figure 3.**
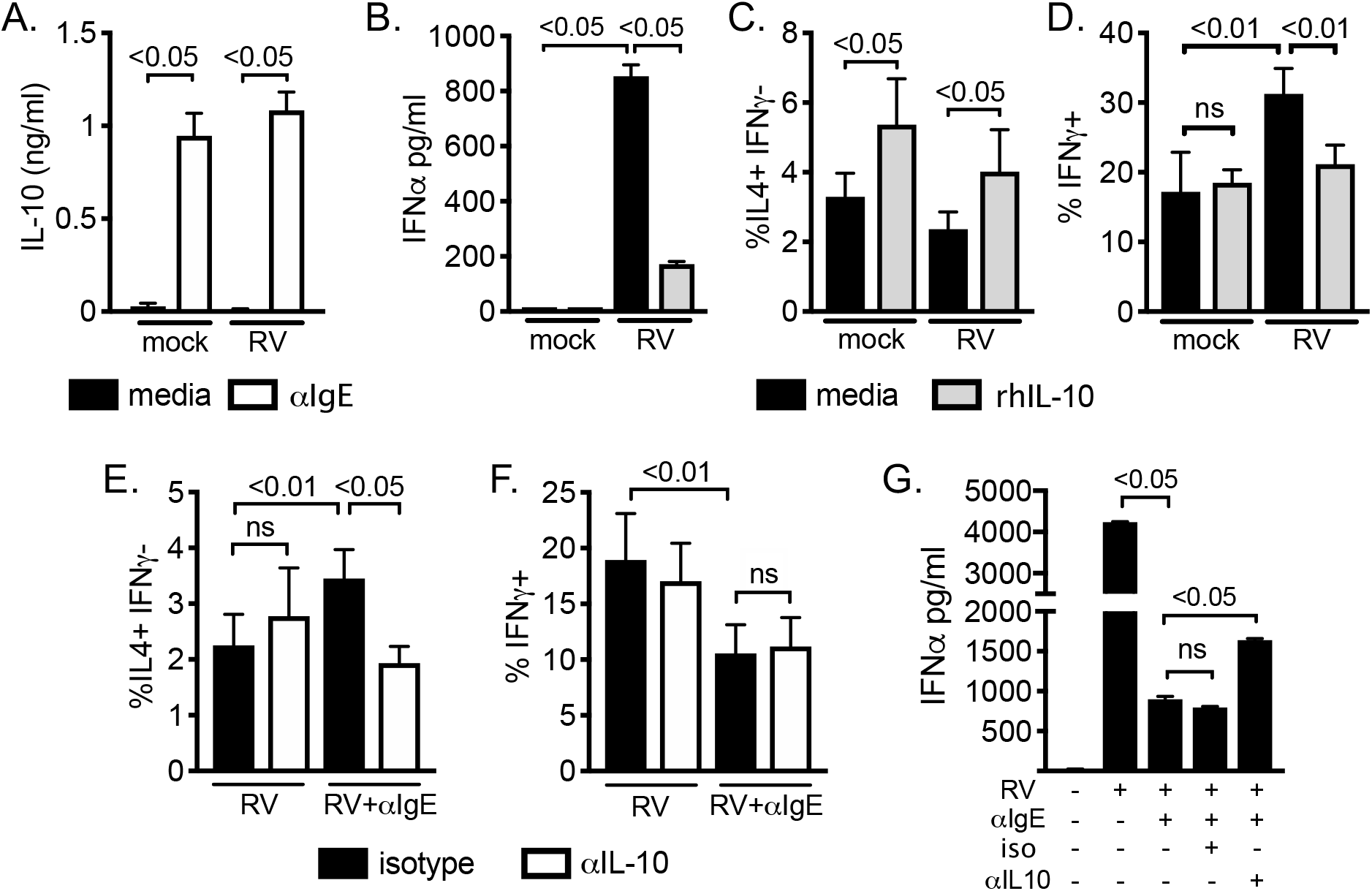
IgE-induced IL-10 is required and sufficient for IgE-mediated enhancement of RV-driven Th2 priming. (A) Primary human monocytes were exposed to mock or RV conditions in the presence or absence of IgE crosslinking antibody, and supernatants harvested for IL-10 ELISA analysis at 24 hours, representative of n=3 experiments shown, p values by one-way ANOVA with post-hoc comparison. (B-D) rhIL-10 (10ng/mL) was added to monocyte cultures in the presence or absence of RV-exposure; (B) IFNα was measured in monocyte supernatants at 24 hours post infection by ELISA analysis, representative of n=3 experiments shown, p values by one-way ANOVA with post-hoc comparison. (C-D) At 24 hours post infection monocytes were co-cultured with allogeneic naïve CD4 T cells to measure either (C) Th2 (%IL-4+IFN*γ*-) or (D) Th1 (%IFN*γ*+) T cell priming. (E-G) IL-10/IL-10Rα blocking (αIL-10) or isotype control antibodies (10μg/mL) were added to RV-exposed monocyte-T cell co-cultures as indicated and T cells evaluated for (E) Th2 or (F) Th1 differentiation. (G) IFNα was measured in co-culture supernatants at day 3 by ELISA, representative of n=3 experiments shown, p values by one-way ANOVA with post-hoc comparison. (C-F) Error bars are SEM, p values by two-way ANOVA with post-hoc comparison, p>0.05 was considered non-significant (ns), n=4 experiments.

## Discussion

Our study shows that IgE-mediated allergic stimulation of monocytes enhances RV-driven Th2 priming of naïve CD4 T cells, which is regulated by IgE-induced IL-10 production and inhibition of virus-induced type I IFN. In our proposed model (Figure 4), the combination of IgE-mediated allergic stimulation and RV exposure leads to enhanced Th2 differentiation via IgE-induced effects on monocytes. In a healthy, non-allergic antiviral response, Th2 differentiation is regulated by viral-induced type I IFN, resulting in a Th1-predominant antiviral response. In the setting of IgE-mediated allergic stimulation, however, IgE-induced IL-10 production inhibits monocyte IFN responses to RV infection. The impaired IFN response combined with other unknown IgE-mediated factors such as altered expression of co-stimulatory molecules, cytokines, chemokines, etc., allows for increased Th2 differentiation and associated allergic inflammation due to the loss of IFN-induced negative regulation on Th2 cell development.

**Figure 4.**
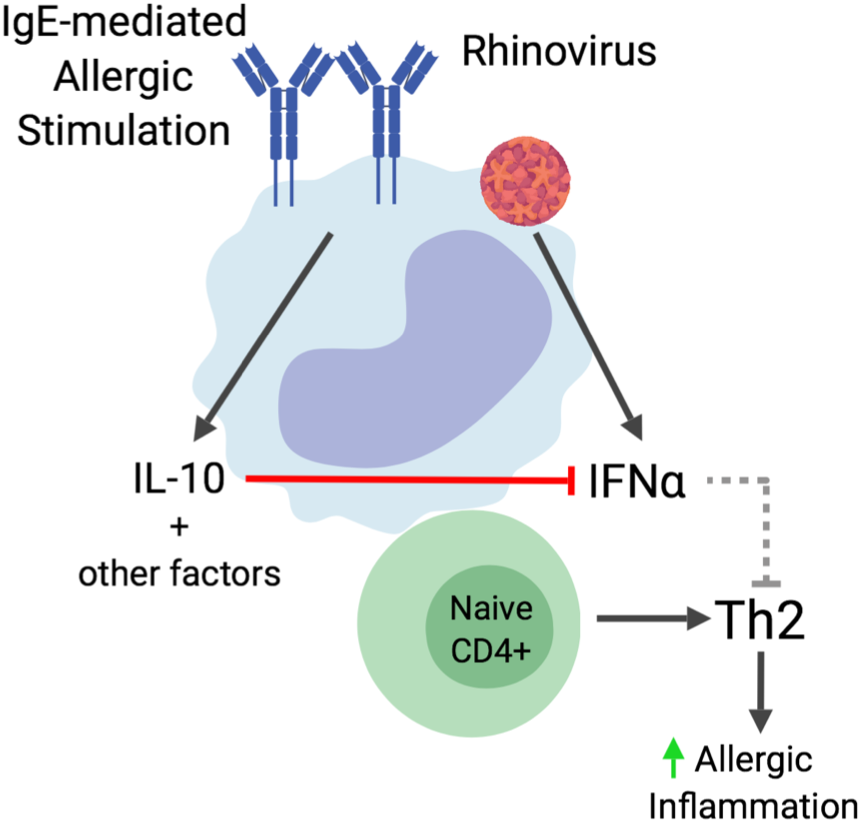
Model of IgE-mediated enhancement of RV-induced monocyte-driven Th2 development. IgE-mediated allergic stimulation combined with RV exposure promotes Th2 priming by monocytes. IgE-induced effects including IL-10 production inhibit RV-driven type I IFN production by monocytes. The reduction of type I IFN-mediated negative regulation of Th2 development combined with other unknown Th2-promoting factors, such as co-stimulatory receptors or cytokines, drive increased Th2 development. This altered antiviral response may contribute to enhanced allergic inflammation.

Although the impact of clinical allergic disease status was not evaluated in our study, we did measure monocyte FcεRIα expression which is known to positively correlate with serum IgE levels and is an established surrogate marker for atopy (32). The correlation between monocyte surface FcεRIα expression and IgE-enhanced RV-driven Th2 differentiation (Fig. 1B) suggests that atopic status may influence IgE-induced antiviral Th2 priming *in vivo*. We did not observe similar correlations with Th2 development by either RV alone (Fig. 1B) or IgE crosslinking in the absence of virus (data not shown), highlighting the relevance of this finding to virus-induced allergic disease. Furthermore, our observation that IgE crosslinking enhanced RV-specific Th2 priming in an autologous model suggests that exposure to relevant allergens in atopic individuals could promote increased RV-induced inflammatory responses during infection. Due to the technical challenges of the autologous co-culture system in measuring virus-specific T cell development, performing additional mechanistic studies were not possible. However, this finding is consistent with results obtained from our allogeneic monocyte-T cell co-culture system and further supports an interaction between allergic stimulation and RV infection in promoting Th2 inflammation. These data have significant implications regarding RV-induced development and exacerbation of allergic diseases. Monocytes recruited to the airway epithelium, the site of both RV infection and aeroallergen exposure, likely encounter allergen-induced IgE crosslinking and virus exposure simultaneously. IgE-mediated local monocyte IL-10 production and inhibition of type I IFN via feedback mechanisms on monocytes and paracrine effects on airway epithelial cells and other APCs could significantly impact local inflammatory responses, potentially enhancing virus replication, virus-induced tissue destruction, and virus-induced Th2 inflammation.

Previous work has primarily focused on Th2 priming in the context of either IgE- or RV-mediated processes alone. Our study begins to address the synergistic role of two signals both epidemiologically linked to allergic disease development and pathogenesis. The differences induced by IgE crosslinking and virus exposure of monocytes, alone versus in concert, demonstrate the importance of evaluating the combined effects of these stimuli on cellular functions. While IgE crosslinking alone promoted Th2 priming, increased Th1 differentiation and T cell proliferation was also observed in our study. Consistent with this, others have reported that IgE-mediated effects on dendritic cells promote antigen-specific Th2 priming and T cell proliferation (33). The IgE-mediated increase in Th1 priming in the absence of virus, while less than virus-induced Th1 differentiation, does support previous work demonstrating a role for Th1 cells in allergic airway inflammation (34, 35). By evaluating Th1 priming in both the presence and absence of RV, we have established a potential role for IgE-mediated inhibition of virus-induced type I IFN in regulating both Th1 and Th2 responses (Fig. 2). Additional investigation, which is beyond the scope of this study, is needed to define specific IgE-mediated factors responsible for these responses.

Deficient antiviral responses, specifically type I IFN, have been repeatedly implicated in the pathogenesis of allergic disease. Enhancement of antiviral IFN responses therefore has a potential therapeutic role and represents an area of intense clinical interest. Multiple approaches to augment antiviral pathways to treat allergic diseases are being investigated (36–38). In a clinical trial of inhaled IFN*β* to prevent virus-induced asthma exacerbations, IFNβ treatment enhanced antiviral gene transcription in sputum cells and in sub-analysis of subjects with severe disease, decreased asthma symptom scores and promoted faster recovery of morning peak expiratory flow (39). In a murine model of RV-induced allergic airway disease, IFNα treatment blocked allergic inflammation induced by exposure to house-dust mite and RV infection in a TLR-7 deficient mouse (40). In our system, complementation of the IgE-induced IFN deficit by exogenous rhIFNα treatment fully reversed IgE-enhanced RV-driven Th2 priming and partially restored impaired virus-induced Th1 responses by monocytes (Fig. 2), supporting its critical role in regulating allergic inflammation and potential value as a therapeutic target in atopic diseases.

The role for IL-10 in allergic disease is more complex(17, 18); it has been shown to play a role in allergen tolerance (41) and virus-induced wheezing (16). IL-10 has a broad range of effector functions on both innate immune and T cells including inhibition of monocyte differentiation, downregulation antigen presenting molecules, and blocking monocyte-driven Th1 cell stimulation (Fig. 3D) (15, 31, 42). Atopic individuals have been shown to have elevated serum IL-10 (43) and increased IL-10 producing monocytes which were capable of promoting Th2 cytokine production in T cells *ex vivo* (17). In our system, IL-10 was both required and sufficient to enhance RV-driven Th2 development, however IL-10 alone did not increase Th2 priming above IL-10 treated monocytes in the absence of virus (Fig. 3D). This is in contrast to our observations comparing IgE crosslinking in the presence and absence of RV-exposure (Fig 1I) where, despite both conditions containing IL-10, Th2 priming by RV combined with IgE crosslinking was greater than IgE crosslinking alone. This could be due to a temporal relationship between IgE-mediated IL-10 production and subsequent inhibition of type I IFN in our system. Alternatively, which we believe is more likely, is the requirement of other, as yet undefined, IgE-mediated effects on monocytes (Fig. 4). The potential contributions of additional IgE-mediated factor(s) is also supported by our finding that although exogenous IL-10, like IgE crosslinking, was capable of suppressing RV-driven Th1 priming (Fig. 3D), IL-10 blockade was not sufficient to reverse IgE-mediated Th1 inhibition (Fig. 3F). This is consistent with previous reports that IL-10 requires additional factors such as antigen presentation, co-stimulation, or other cytokines to modulate APC-driven T cell differentiation (29, 30). We previously showed that IgE crosslinking inhibits influenza-induced upregulation of antigen presenting molecules. In that system, blocking IL-10 was similarly insufficient to reverse IgE-mediated inhibition of influenza-driven Th1 priming (12). Thus, additional IgE-induced effects such as modulation of monocyte antigen-presenting molecules, expression of co-stimulatory molecules that enhance Th2 priming (29, 44), or other cytokines (12, 13, 45), combined with IL-10 production and the loss of antiviral IFN could also regulate IgE-mediated Th1 and Th2 development. While our data demonstrate a cellular mechanism by which IgE-induced IL-10 inhibits antiviral IFN responses to promote virus-induced Th2 priming, additional investigation is needed to identify the specific cell signaling events involved. Further studies to elucidate the complex interaction between IgE-mediated signals and RV-induced monocyte-driven T cell development are likely to identify additional factors critical in these processes.

To our knowledge, this is the first report that IgE-mediated stimulation promotes RV-induced Th2 priming by human monocytes and that impaired monocyte IFN responses may specifically contribute to Th2-mediated inflammation. While IL-10 has been shown to inhibit virus-induced type I IFN responses in human PBMCs (46), our results support a previously undescribed role for IL-10 in the inhibition of virus-induced IFN responses to promote Th2 inflammation. Beyond allergic disease, these findings highlight a potential regulatory mechanism by which IgE-mediated pathways may have evolved to regulate Th1-driven inflammation in favor of Th2 responses - *eg.* helminth infections or promoting antigen tolerance. Finally, we demonstrate that IFNα can reverse IgE-mediated induction of antiviral Th2 differentiation in human cells. Given the close link between RV infection and allergic disease development and exacerbation, and the critical role of monocytes in airway immune responses, this study provides another potential mechanism by which IgE-mediated activation synergizes with RV infection to drive allergic inflammation. Identification of additional molecular mechanisms underlying these processes could lead to potential therapeutic targets for improved treatment and prevention of viral exacerbations of allergic diseases.

## Methods

### Human Subjects

Samples were acquired from 2 sources: leukocyte-enriched blood samples from unknown donors through a local blood bank and blood samples from healthy volunteers. Unknown donors were deemed healthy enough for routine blood donation by the local blood bank, no information on atopic status was obtained from these unknown donors. Studies were approved by the UT Southwestern institutional review board (Study # STU 122010-139). For known volunteers, written informed consent and assent were obtained.

### Reagents and Media

Phosphate Buffered Saline (PBS) with fetal bovine serum (FBS) and EDTA (complete PBS, cPBS) and complete Roswell Park Memorial Institute Medium 1640 (cRPMI) were prepared as previously described (13). Rabbit anti-human IgE (αIgE) was purchased from Bethyl Laboratories (Montgomery, TX) and rabbit whole IgG control antibody (IgG) from Jackson ImmunoResearch (Westgrove, PA). Unless specified, recombinant human (rh) cytokines, anti-human cytokine and receptor antibodies were purchased from R&D Systems (Minneapolis, MN).

### Purification of Immune Cells from Blood

Monocytes and naïve CD4^+^ T cells were purified as previously described (12, 13). Purity was assessed by flow cytometry; monocytes were identified as CD14+ and naïve CD4^+^ T cells as CD3+CD4+CD45RA+. Samples with <85% purity were excluded.

### Monocyte Culture Conditions

Monocytes were cultured in cRPMI with recombinant human (rh) M-CSF (1 ng/ml) at a concentration of 1 x 10^6^ cells/ml. αIgE (10 μg/ml), rabbit IgG isotype control (10 μg/ml), or IL-10 (10 ng/ml), anti-IL-10 and anti-IL-10 receptor (IL-10Rα) (5μg/mL each), or mouse IgG isotype controls were added to monocyte cultures at the start of culture. Purified Rhinovirus A serotype 16 (RV) (kindly provided by Jim Gern and Yury Bochkov, U Wisconsin) was added at a multiplicity of infection (MOI) 10. Cultures were incubated at 37°C for indicated times. Cells and supernatants were harvested for analysis or for T cell co-culture experiments.

### Monocyte-T Cell Co-Cultures

Monocytes were cultured as above in the presence or absence of αIgE, control IgG, or RV at 37°C for 18-24 hours and then mixed 1:1 with allogeneic naïve CD4^+^ T cells. Cells were cultured in media containing 50 international units per mL (IU/mL) rhIL-2. For select experiments, mouse anti-human antibodies against IL-10, IL-10Rα, or IFNAR and IgG1 or IgG2b isotype controls were added at 5-10μg/mL each for a total mouse IgG concentration of 10 μg/ml prior to co-culture with T cells. As indicated, recombinant human (rh) IFNα2a (1000-10,000 IU/mL) was added at the initiation of co-culture. After 3 days of co-culture, T cells were expanded 1:5 in media with IL-2 and cultured for an additional 3-4 days. Cells were then rested in media without IL-2 overnight and restimulated for flow cytometry analysis as previously described (12). In a subset of experiments to evaluate cell proliferation, T cells were labeled prior to culture with the cell proliferation dye, VPD450, per manufacturer’s instructions (BD Biosciences, San Jose).

To evaluate virus-specific monocyte-driven Th2 development, autologous PBMCs were isolated from known donors, monocytes and naïve CD4 T cells purified as above and co-cultures were performed. Monocytes were either mock treated (media alone) or exposed as above to RV with and without IgE crosslinking or isotype IgG control. At 24 hours, monocytes were co-cultured at 1:1 ratio with autologous naïve CD4 T cells in U-bottom 96-well plates. Cells were co-cultured for 12 days in the presence of rhIL-2. On days 6 and 9, 50μl of media was removed and replaced with fresh media containing 10 IU/mL IL-2. On day 12, cells were expanded to 1:10 vol:vol (2mL) into 24-well plates in media with 15 IU/mL IL-2 for an additional 4 days. Like conditions were then combined, media removed, and cells rested in 6-well dishes in media without IL-2 overnight prior to restimulation. Two days prior to T cell restimulation, autologous PBMCs frozen at the initial blood draw were thawed and exposed to RV at MOI=5 for 48 hours. Just prior to restimulation of primed T cells, PBMCs were x-ray irradiated with a dose of 13Gy (450 seconds at 250kV potential, 15mA, XRAD 320 model). Irradiated PBMCs were counted and added to the primed naïve CD4 T cells at a 1:1 ratio and cultured for 24 hours, with monensin (golgi block) added in the final 6 hours. Cells were fixed in 2% paraformaldehyde and stained for flow cytometry analysis with the following antibodies: CD4 APC, HLA-DR FITC, IL-4 PE, and IFN*γ* PE-Cy7. A minimum of 40,000 events were acquired and primed T cells were identified as CD4+ HLA-DR low/intermediate and then quantified based on their cytokine expression (IL-4 or IFN*γ* single positive). Irradiated PBMCs were cultured alone to control for residual cytokine expression following irradiation. We confirmed little to no detectable cytokine production by irradiated PBMCs (data not shown). Cytokine expressing cells were normalized as the total number of cells per 1000 CD4+ events.

### Flow Cytometry Analysis of Surface Antigens

The following fluorochrome-conjugated anti-human antibodies or molecules were used: CD3-peridinin chlorophyll protein complex (PerCP), CD4-allophycocyanin (APC), CD14-V450, CD45RA-phycoethrin (PE), FcεRIα-fluorescein (FITC), HLA-DR-APC-Cy7, and HLA-DR-FITC (BD Pharmingen, San Diego, CA). Staining and flow cytometry were performed as previously described (13). Samples were acquired on a BD LSR II flow cytometer (BD Biosciences) and analyzed with the FlowJo 8 software (FLOWJO, LLC, Ashland, OR).

### Flow Cytometry Analysis of Intracellular Cytokines

The following antibodies were used: IFNγ-PE-Cy7 and IL-4-PE (BD Biosciences, San Jose, CA). Cells were fixed in 2% paraformaldehyde and permeablized with 0.1% saponin in PBS to detect intracellular antigens. Samples were acquired and analyzed as above described. Gating of cytokine-expressing cells was guided by staining negative control (unstimulated cells) and positive control T cells differentiated under Th1- or Th2-inducing conditions (23). Percentages of total cytokine-positive populations were then determined. Th2 cells were identified as IL-4 single positive; Th1 cells were identified as the total IFN*γ* population (Supp. Fig. 1). To measure cell proliferation, divided cells were identified as VPD450 low, as compared to unstimulated naïve T cells.

### Cytokine Secretion Analysis

Supernatants were stored at −80°C until use. The following ELISA kits were used according to manufacturer recommendations: ELISA Max human ELISA for IL-10 (Biolegend, San Diego, CA), anti-pan human IFN alpha (Interferon Source, Cincinnati, OH). Absorbance was measured on a Biorad iMark microplate reader (Biorad, Hercules, CA) per manufacturer’s instructions.

### Data Analysis and Statistics

Data are presented as means ± standard error of the mean (SEM) or standard deviation (SD). In experiments containing 3 or more conditions, one-way or two-way (for experiments with multiple donors) repeated measures ANOVA and pairwise Tukey’s post hoc comparisons were performed. For experiments comparing 2 conditions, paired t tests were performed with Holm-Sidak correction where appropriate. p<0.05 was considered significant, pertinent p values are noted in figures; significant as <0.05, <0.01, and <0.001, or >0.05 for non-significant (ns) values. All statistical analyses were performed using GraphPad Prism version 8.

## Abbreviations

APC: antigen presenting cell
αIgE: Rabbit anti-human IgE antibody
FcεRI: high-affinity IgE receptor
IFN: interferon
IgG: Rabbit whole IgG isotype control antibody
IL: interleukin
PBMCs: peripheral blood mononuclear cells
pDC: plasmacytoid dendritic cells
rh: recombinant human
RV: Rhinovirus
Th: CD4 helper T cell
TLR: toll-like receptor

## Acknowledgements

We thank the participants, without whom this study would not be possible. We thank Jim Gern, M.D. and Yury Bochkov, Ph.D. (U Wisconsin) for providing highly purified RV-16 virus. We also acknowledge Yaging Gao and Robert Maples specifically for assistance with PBMC irradiation, the members of the Farrar lab (UTSW) for helpful discussions and suggestions, and Angela Mobley and staff of the UTSW Immunology Flow Cytometry core for assistance with flow cytometry analysis. Figure 4 was created with Biorender.

## Figure Legends

**Supplemental Figure 1. IgE crosslinking enhances RV-induced monocyte driven Th2 differentiation.** Flow cytometry plots of a representative experiment from Figure 1A are shown. Primary human monocytes were exposed to RV in the presence or absence of IgE crosslinking (αIgE) or isotype control (IgG) antibody. Monocytes were co-cultured with allogeneic naïve CD4 T cells to evaluate monocyte-induced T cell development. Cells were gated based on cytokine expression as a % of total cells in the live, single cell gate. Th2 cells were identified as IL-4 single positive while Th1 cells (Supp. Fig. 2) were identified as the total IFN*γ* positive population.

**Supplemental Figure 2. IgE crosslinking impairs RV-driven Th1 development.** Monocytes were co-cultured with allogeneic naïve CD4 T cells and Th1 priming was evaluated as determined by % IFN*γ* positive population. (A) RV-driven Th1 priming is inhibited by IgE crosslinking, n=17 donor pairs, while (B) IgE crosslinking alone slightly enhances Th1 development, n=15 donor pairs. Violin plots show mean depicted as black line with individual experiments as open circles. p values obtained by two-way ANOVA with post-hoc comparison.

**Supplemental Figure 3. Type I interferon regulates monocyte-driven T cell development in monocyte-T cell co-cultures.** Primary human monocytes were co-cultured with allogeneic naïve CD4 T cells +/- 1000IU/mL rhIFNα2a (IFNα). T cells were then evaluated for either (A) Th2 or (B) Th1 differentiation. p values by two-way ANOVA with post-hoc analysis, n=3 experiments.

## References

1. Johnston, N. W., S. L. Johnston, J. M. Duncan, J. M. Greene, T. Kebadze, P. K. Keith, M. Roy, S. Waserman, and M. R. Sears. 2005. The September epidemic of asthma exacerbations in children: a search for etiology. J Allergy Clin Immunol 115: 132–138.

2. Khetsuriani, N., N. N. Kazerouni, D. D. Erdman, X. Lu, S. C. Redd, L. J. Anderson, and W. G. Teague. 2007. Prevalence of viral respiratory tract infections in children with asthma. J Allergy Clin Immunol 119: 314–321.

3. Rubner, F. J., D. J. Jackson, M. D. Evans, R. E. Gangnon, C. J. Tisler, T. E. Pappas, J. E. Gern, and R. F. Lemanske, Jr. 2016. Early life rhinovirus wheezing, allergic sensitization, and asthma risk at adolescence. J Allergy Clin Immunol.

4. Zambrano, J. C., H. T. Carper, G. P. Rakes, J. Patrie, D. D. Murphy, T. A. Platts-Mills, F. G. Hayden, J. M. Gwaltney, Jr., T. K. Hatley, A. M. Owens, and P. W. Heymann. 2003. Experimental rhinovirus challenges in adults with mild asthma: response to infection in relation to IgE. J Allergy Clin Immunol 111: 1008–1016.

5. Teach, S. J., M. A. Gill, A. Togias, C. A. Sorkness, S. J. Arbes, Jr., A. Calatroni, J. J. Wildfire, P. J. Gergen, R. T. Cohen, J. A. Pongracic, C. M. Kercsmar, G. K. Khurana Hershey, R. S. Gruchalla, A. H. Liu, E. M. Zoratti, M. Kattan, K. A. Grindle, J. E. Gern, W. W. Busse, and S. J. Szefler. 2015. Preseasonal treatment with either omalizumab or an inhaled corticosteroid boost to prevent fall asthma exacerbations. J Allergy Clin Immunol 136: 1476–1485.

6. Casale, T. B., J. Condemi, C. LaForce, A. Nayak, M. Rowe, M. Watrous, M. McAlary, A. Fowler-Taylor, A. Racine, N. Gupta, R. Fick, and G. Della Cioppa. 2001. Effect of omalizumab on symptoms of seasonal allergic rhinitis: a randomized controlled trial. JAMA : the journal of the American Medical Association 286: 2956–2967.

7. Kim, D. H., K. Y. Park, B. J. Kim, M. N. Kim, and S. K. Mun. 2013. Anti-immunoglobulin E in the treatment of refractory atopic dermatitis. Clinical and experimental dermatology 38: 496–500.

8. Esquivel, A., W. W. Busse, A. Calatroni, A. G. Togias, K. G. Grindle, Y. A. Bochkov, R. S. Gruchalla, M. Kattan, C. M. Kercsmar, G. Khurana Hershey, H. Kim, P. Lebeau, A. H. Liu, S. J. Szefler, S. J. Teach, J. B. West, J. Wildfire, J. A. Pongracic, and J. E. Gern. 2017. Effects of Omalizumab on Rhinovirus Infections, Illnesses and Exacerbations of Asthma. American journal of respiratory and critical care medicine.

9. Zhu, J., S. D. Message, Y. Qiu, P. Mallia, T. Kebadze, M. Contoli, C. K. Ward, E. S. Barnathan, M. A. Mascelli, O. M. Kon, A. Papi, L. A. Stanciu, P. K. Jeffery, and S. L. Johnston. 2014. Airway inflammation and illness severity in response to experimental rhinovirus infection in asthma. Chest 145: 1219–1229.

10. Gill, M. A., K. Long, T. Kwon, L. Muniz, A. Mejias, J. Connolly, L. Roy, J. Banchereau, and O. Ramilo. 2008. Differential recruitment of dendritic cells and monocytes to respiratory mucosal sites in children with influenza virus or respiratory syncytial virus infection. J Infect Dis 198: 1667–1676.

11. Eguiluz-Gracia, I., A. Bosco, R. Dollner, G. R. Melum, M. H. Lexberg, A. C. Jones, S. A. Dheyauldeen, P. G. Holt, E. S. Baekkevold, and F. L. Jahnsen. 2016. Rapid recruitment of CD14(+) monocytes in experimentally induced allergic rhinitis in human subjects. J Allergy Clin Immunol 137: 1872–1881 e1812.

12. Rowe, R. K., D. M. Pyle, A. R. Tomlinson, T. Lv, Z. Hu, and M. A. Gill. 2017. IgE cross-linking impairs monocyte antiviral responses and inhibits influenza-driven TH1 differentiation. J Allergy Clin Immunol.

13. Pyle, D. M., V. S. Yang, R. S. Gruchalla, J. D. Farrar, and M. A. Gill. 2013. IgE cross-linking critically impairs human monocyte function by blocking phagocytosis. J Allergy Clin Immunol 131: 491–500 e495.

14. Novak, N., W. M. Peng, T. Bieber, and C. Akdis. 2013. FcepsilonRI stimulation promotes the differentiation of histamine receptor 1-expressing inflammatory macrophages. Allergy 68: 454–461.

15. Novak, N., T. Bieber, and N. Katoh. 2001. Engagement of Fc epsilon RI on human monocytes induces the production of IL-10 and prevents their differentiation in dendritic cells. J Immunol 167: 797–804.

16. Bont, L., C. J. Heijnen, A. Kavelaars, W. M. van Aalderen, F. Brus, J. T. Draaisma, S. M. Geelen, and J. L. Kimpen. 2000. Monocyte IL-10 production during respiratory syncytial virus bronchiolitis is associated with recurrent wheezing in a one-year follow-up study. American journal of respiratory and critical care medicine 161: 1518–1523.

17. Prasse, A., M. Germann, D. V. Pechkovsky, A. Markert, T. Verres, M. Stahl, I. Melchers, W. Luttmann, J. Muller-Quernheim, and G. Zissel. 2007. IL-10-producing monocytes differentiate to alternatively activated macrophages and are increased in atopic patients. J Allergy Clin Immunol 119: 464–471.

18. Raedler, D., S. Illi, L. A. Pinto, E. von Mutius, T. Illig, M. Kabesch, and B. Schaub. 2013. IL10 polymorphisms influence neonatal immune responses, atopic dermatitis, and wheeze at age 3 years. J Allergy Clin Immunol 131: 789–796.

19. Jackson, D. J., H. Makrinioti, B. M. Rana, B. W. Shamji, M. B. Trujillo-Torralbo, J. Footitt, D.-R. Jerico, A. G. Telcian, A. Nikonova, J. Zhu, J. Aniscenko, L. Gogsadze, E. Bakhsoliani, S. Traub, J. Dhariwal, J. Porter, D. Hunt, T. Hunt, T. Hunt, L. A. Stanciu, M. Khaitov, N. W. Bartlett, M. R. Edwards, O. M. Kon, P. Mallia, N. G. Papadopoulos, C. A. Akdis, J. Westwick, M. J. Edwards, D. J. Cousins, R. P. Walton, and S. L. Johnston. 2014. IL-33-dependent type 2 inflammation during rhinovirus-induced asthma exacerbations in vivo. American journal of respiratory and critical care medicine 190: 1373–1382.

20. Schroeder, J. T., A. P. Bieneman, K. L. Chichester, R. G. Hamilton, H. Xiao, S. S. Saini, and M. C. Liu. 2010. Decreases in human dendritic cell-dependent T(H)2-like responses after acute in vivo IgE neutralization. J Allergy Clin Immunol 125: 896–901 e896.

21. Bellinghausen, I., U. Brand, J. Knop, and J. Saloga. 2000. Comparison of allergen-stimulated dendritic cells from atopic and nonatopic donors dissecting their effect on autologous naive and memory T helper cells of such donors. J Allergy Clin Immunol 105: 988–996.

22. Huber, J. P., and J. D. Farrar. 2011. Regulation of effector and memory T-cell functions by type I interferon. Immunology 132: 466–474.

23. Huber, J. P., H. J. Ramos, M. A. Gill, and J. D. Farrar. 2010. Cutting edge: Type I IFN reverses human Th2 commitment and stability by suppressing GATA3. J Immunol 185: 813–817.

24. Durrani, S. R., D. J. Montville, A. S. Pratt, S. Sahu, M. K. DeVries, V. Rajamanickam, R. E. Gangnon, M. A. Gill, J. E. Gern, R. F. Lemanske, Jr., and D. J. Jackson. 2012. Innate immune responses to rhinovirus are reduced by the high-affinity IgE receptor in allergic asthmatic children. J Allergy Clin Immunol 130: 489–495.

25. Gill, M. A., A. H. Liu, A. Calatroni, R. Z. Krouse, B. Shao, A. Schiltz, J. E. Gern, A. Togias, and W. W. Busse. 2017. Enhanced plasmacytoid dendritic cell antiviral responses after omalizumab. J Allergy Clin Immunol.

26. Gill, M. A., G. Bajwa, T. A. George, C. C. Dong, Dougherty, II, N. Jiang, V. N. Gan, and R. S. Gruchalla. 2010. Counterregulation between the FcepsilonRI pathway and antiviral responses in human plasmacytoid dendritic cells. J Immunol 184: 5999–6006.

27. Longhi, M. P., C. Trumpfheller, J. Idoyaga, M. Caskey, I. Matos, C. Kluger, A. M. Salazar, M. Colonna, and R. M. Steinman. 2009. Dendritic cells require a systemic type I interferon response to mature and induce CD4+ Th1 immunity with poly IC as adjuvant. J Exp Med 206: 1589–1602.

28. Santini, S. M., C. Lapenta, S. Donati, F. Spadaro, F. Belardelli, and M. Ferrantini. 2011. Interferon-alpha-conditioned human monocytes combine a Th1-orienting attitude with the induction of autologous Th17 responses: role of IL-23 and IL-12. PLoS One 6: e17364.

29. Liu, L., B. E. Rich, J. Inobe, W. Chen, and H. L. Weiner. 1998. Induction of Th2 cell differentiation in the primary immune response: dendritic cells isolated from adherent cell culture treated with IL-10 prime naive CD4+ T cells to secrete IL-4. Int Immunol 10: 1017–1026.

30. Hsieh, C. S., A. B. Heimberger, J. S. Gold, A. O’Garra, and K. M. Murphy. 1992. Differential regulation of T helper phenotype development by interleukins 4 and 10 in an alpha beta T-cell-receptor transgenic system. Proceedings of the National Academy of Sciences of the United States of America 89: 6065–6069.

31. Fiorentino, D. F., A. Zlotnik, P. Vieira, T. R. Mosmann, M. Howard, K. W. Moore, and A. O’Garra. 1991. IL-10 acts on the antigen-presenting cell to inhibit cytokine production by Th1 cells. J Immunol 146: 3444–3451.

32. Sihra, B. S., O. M. Kon, J. A. Grant, and A. B. Kay. 1997. Expression of high-affinity IgE receptors (Fc epsilon RI) on peripheral blood basophils, monocytes, and eosinophils in atopic and nonatopic subjects: relationship to total serum IgE concentrations. J Allergy Clin Immunol 99: 699–706.

33. Sallmann, E., B. Reininger, S. Brandt, N. Duschek, E. Hoflehner, E. Garner-Spitzer, B. Platzer, E. Dehlink, M. Hammer, M. Holcmann, H. C. Oettgen, U. Wiedermann, M. Sibilia, E. Fiebiger, A. Rot, and D. Maurer. 2011. High-affinity IgE receptors on dendritic cells exacerbate Th2-dependent inflammation. J Immunol 187: 164–171.

34. Hansen, G., G. Berry, R. H. DeKruyff, and D. T. Umetsu. 1999. Allergen-specific Th1 cells fail to counterbalance Th2 cell-induced airway hyperreactivity but cause severe airway inflammation. J Clin Invest 103: 175–183.

35. Wisniewski, J. A., L. M. Muehling, J. D. Eccles, B. J. Capaldo, R. Agrawal, D. A. Shirley, J. T. Patrie, L. J. Workman, A. J. Schuyler, M. G. Lawrence, W. G. Teague, and J. A. Woodfolk. 2018. TH1 signatures are present in the lower airways of children with severe asthma, regardless of allergic status. J Allergy Clin Immunol 141: 2048–2060 e2013.

36. Casale, T. B., J. Cole, E. Beck, C. F. Vogelmeier, J. Willers, C. Lassen, A. Hammann-Haenni, L. Trokan, P. Saudan, and M. E. Wechsler. 2015. CYT003, a TLR9 agonist, in persistent allergic asthma - a randomized placebo-controlled Phase 2b study. Allergy 70: 1160–1168.

37. Greiff, L., C. Ahlstrom-Emanuelsson, M. Alenas, G. Almqvist, M. Andersson, A. Cervin, J. Dolata, S. Lindgren, A. Martensson, B. Young, and H. Widegren. 2015. Biological effects and clinical efficacy of a topical Toll-like receptor 7 agonist in seasonal allergic rhinitis: a parallel group controlled phase IIa study. Inflamm Res 64: 903–915.

38. Silkoff, P. E., S. Flavin, R. Gordon, M. J. Loza, P. J. Sterk, R. Lutter, Z. Diamant, R. B. Turner, B. J. Lipworth, D. Proud, D. Singh, A. Eich, V. Backer, J. E. Gern, C. Herzmann, S. A. Halperin, T. T. Mensinga, A. M. Del Vecchio, P. Branigan, L. San Mateo, F. Baribaud, E. S. Barnathan, and S. L. Johnston. 2018. Toll-like receptor 3 blockade in rhinovirus-induced experimental asthma exacerbations: A randomized controlled study. J Allergy Clin Immunol 141: 1220–1230.

39. Djukanovic, R., T. Harrison, S. L. Johnston, F. Gabbay, P. Wark, N. C. Thomson, R. Niven, D. Singh, H. K. Reddel, D. E. Davies, R. Marsden, C. Boxall, S. Dudley, V. Plagnol, S. T. Holgate, P. Monk, and I. S. Group. 2014. The effect of inhaled IFN-beta on worsening of asthma symptoms caused by viral infections. A randomized trial. American journal of respiratory and critical care medicine 190: 145–154.

40. Hatchwell, L., A. Collison, J. Girkin, K. Parsons, J. Li, J. Zhang, S. Phipps, D. Knight, N. W. Bartlett, S. L. Johnston, P. S. Foster, P. A. Wark, and J. Mattes. 2015. Toll-like receptor 7 governs interferon and inflammatory responses to rhinovirus and is suppressed by IL-5-induced lung eosinophilia. Thorax 70: 854–861.

41. Chung, F. 2001. Anti-inflammatory cytokines in asthma and allergy: interleukin-10, interleukin-12, interferon-gamma. Mediators Inflamm 10: 51–59.

42. de Waal Malefyt, R., J. Haanen, H. Spits, M. G. Roncarolo, A. te Velde, C. Figdor, K. Johnson, R. Kastelein, H. Yssel, and J. E. de Vries. 1991. Interleukin 10 (IL-10) and viral IL-10 strongly reduce antigen-specific human T cell proliferation by diminishing the antigen-presenting capacity of monocytes via downregulation of class II major histocompatibility complex expression. J Exp Med 174: 915–924.

43. Wong, C. K., C. Y. Ho, F. W. Ko, C. H. Chan, A. S. Ho, D. S. Hui, and C. W. Lam. 2001. Proinflammatory cytokines (IL-17, IL-6, IL-18 and IL-12) and Th cytokines (IFN-gamma, IL-4, IL-10 and IL-13) in patients with allergic asthma. Clin Exp Immunol 125: 177–183.

44. Rogers, P. R., and M. Croft. 2000. CD28, Ox-40, LFA-1, and CD4 modulation of Th1/Th2 differentiation is directly dependent on the dose of antigen. J Immunol 164: 2955–2963.

45. Heijink, I. H., E. Vellenga, P. Borger, D. S. Postma, J. G. de Monchy, and H. F. Kauffman. 2002. Interleukin-6 promotes the production of interleukin-4 and interleukin-5 by interleukin-2-dependent and -independent mechanisms in freshly isolated human T cells. Immunology 107: 316–324.

46. Payvandi, F., S. Amrute, and P. Fitzgerald-Bocarsly. 1998. Exogenous and endogenous IL-10 regulate IFN-alpha production by peripheral blood mononuclear cells in response to viral stimulation. J Immunol 160: 5861–5868.

