## Supplementary figures and images for "IgE-mediated regulation of IL-10 and type I interferon enhances rhinovirus-induced Th2 priming by primary human monocytes"

### Supplemental Figures

#### Supplemental Figure 1

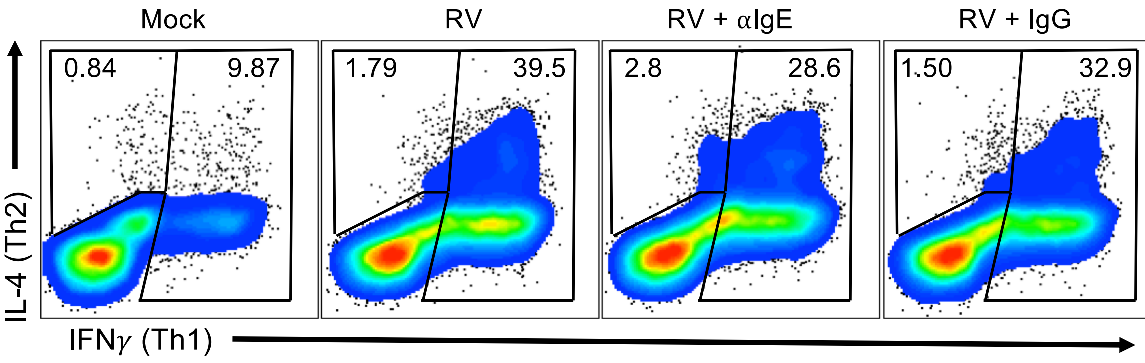

#### Supplemental Figure 2

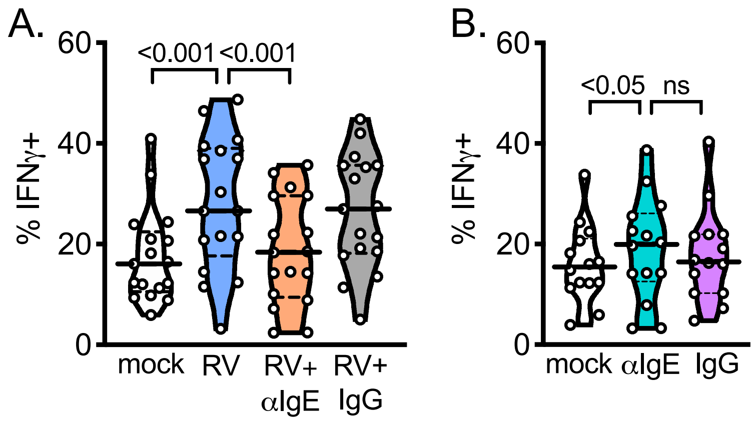

#### Supplemental Figure 3

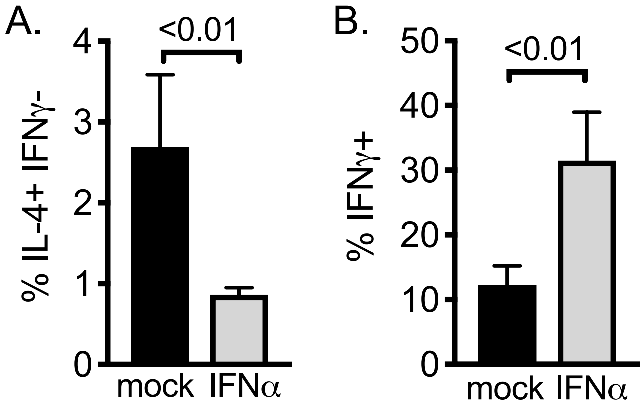
